# A rigid body framework for multi-cellular modelling

**DOI:** 10.1101/2021.02.10.430170

**Authors:** Phillip J. Brown, J. Edward F. Green, Benjamin J. Binder, James M. Osborne

## Abstract

Multi-cellular modelling, where tissues and organs are represented by a collection of individual interacting agents, is a well established field, encapsulating several different approaches. In particular, off-lattice models, which represent cells using points that are free to move in space, have been applied to numerous biological problems in both two and three dimensions. The fact that a cell can be represented by point objects is useful in a wide range of settings, particularly when large populations are involved. However, a purely point-based representation is not naturally equipped to deal with objects that inherently have length like cell boundaries or external membranes.

In this paper we introduce a novel off-lattice modelling framework that exploits rigid body mechanics to represent cells using a collection of one-dimensional edges (rather than zero-dimensional points) in a viscosity-dominated system. The rigid body framework can be used, among other things, to represent cells as free moving polygons, to allow epithelial layers to smoothly interact with themselves, and to model rod shaped cells like bacteria. We demonstrate the value of our new framework by using it in these three applications, showing that this approach can replicate established results, as well as offer solutions to problems that limit the scope of current off-lattice multi-cellular models.

## Main

Multi-cellular modelling is a technique for representing tissues as a collection of individual agents. It can be used to investigate anything from embryogenesis [7] to crypt homeostasis [31]. The agents can be represented by objects on a fixed lattice, or by an “off-lattice” approach using points that are free to move in space. In 1981, Odell et al. [22] demonstrated the ability of off-lattice approaches to investigate morphogenesis caused by the mechanics of cell-cell interactions. Since then they have been used in a vast array of modelling problems and developed into several unique branches. Among them are centre based models where a cell is represented by a single point (for instance, over-lapping spheres [9] and tessellation models [16]), and multi-node models (like the vertex model [20], the sub-cellular element method [21], and the immersed boundary method [27]) where cells are represented by a collection of points. These methods, in a broad sense, are built in a “node-based” framework, since they all calculate motion from interactions between nodes.

Motion in a node-based model is routinely realised mechanically ^2^. The net force **F**_*i*_ applied to each node *i* is calculated, then its movement is determined via equations of motion. Often in cell-based modelling, inertial forces are assumed to be negligible compared to viscous drag forces [24], meaning the motion can be described by the first order differential equations

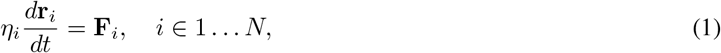

where **r**_*i*_ is the position of the node and *η*_*i*_ its drag coefficient. This is solved numerically for each node to determine its new position in space, often using the Forward Euler method [24]. The net force **F**_*i*_ can be determined either by directly calculating forces (perhaps via a spring law) or by energy methods where the force is the result of energy potentials. Depending on the model, other steps come after this to re-position the nodes, for instance with cell division and death. Notably, the vertex model performs various kinds of *swaps* to change which nodes are part of which cells to allow cells to move through a tissue.

The mathematical simplicity of these approaches make them very popular, and they can achieve a large range of realistic behaviours. Some major modelling works include Odell et. al [22] who first used off-lattice modelling to investigate morphogenesis, Nagai et al. [20] who introduce the common vertex model, Van Leeuwen et al. [31] who make a two-dimensional multi-scale model of the crypt, Osborne et al. [24] who present a thorough comparison of the five predominant models and Okuda et al. [23] who combine a vertex model with a reaction-diffusion system to investigate morphogenesis in three-dimensions. These models often produce “biologically pleasing” images of close-packed cells that show similarities to experiments. They can also capture growth or movement behaviours exhibited at the tissue level.

However, at a fundamental level, a node-based framework is insufficient for modelling barrier objects that separate one region from another, such as a cell surface or an external membrane. A collection of nodes forms the foundation for representing a barrier, but it is the edges joining them together that give a barrier its substance. Hence, to properly represent a barrier, edges need to respond to their environment.

In this paper, we introduce a rigid body modelling framework where cells are modelled by a collection of edges rather than a collection of nodes. Critically, these edges can interact with their simulated environment (e.g. other cells) through sensible forces. We explore several applications of the rigid body framework and demonstrate how they can reproduce established results, and overcome existing issues. These applications are: Bacterial cells modelled by rods growing in a restricted domain; cells modelled as separated polygons used to model a tumour spheroid, and; a contiguous ring of rectangular cells used to represent an epithelial monolayer. Finally, we discuss the power of the framework, its potential applications in cell-based modelling, and highlight areas for future development. A detailed description of the underlying theory is provided in *Methods* and the *Supplementary Material*.

## Results

### A rigid body framework for multi-cellular modelling

The fundamental unit of our modelling framework is a one-dimensional edge. This means that, unlike existing modelling approaches that use zero-dimensional points, we need to consider *where* on the edge forces are applied as well as how the edge *rotates*.

We define an edge by a linked pair of nodes which specify its endpoints, and a line joining the endpoints which represents its interior. Forces can be applied directly to the endpoints, or at some point along the interior, and these forces will generally be caused by interactions with other edges in the model. We limit our scope to only consider interactions amongst endpoints (which we term “node-node” interactions), and between endpoints and interiors (“node-edge” interactions). It is possible to specify interactions between interiors (“edge-edge”), but we leave this for future development.

Figure 1A-C shows how we determine where interaction forces are applied. A circle is centred on an endpoint of an adjacent edge, and its radius expanded until it touches the edge of interest. If the contact point is also an endpoint, we calculate a node-node interaction, while if it is an interior point, we calculate a node-edge interaction. If we use multiple joined edges to construct a cell or a surface, the node may end up interacting with a shared endpoint if the cell is locally convex, or with both edges if the cell is locally concave (Figure 1B and C).

**Figure 1:**
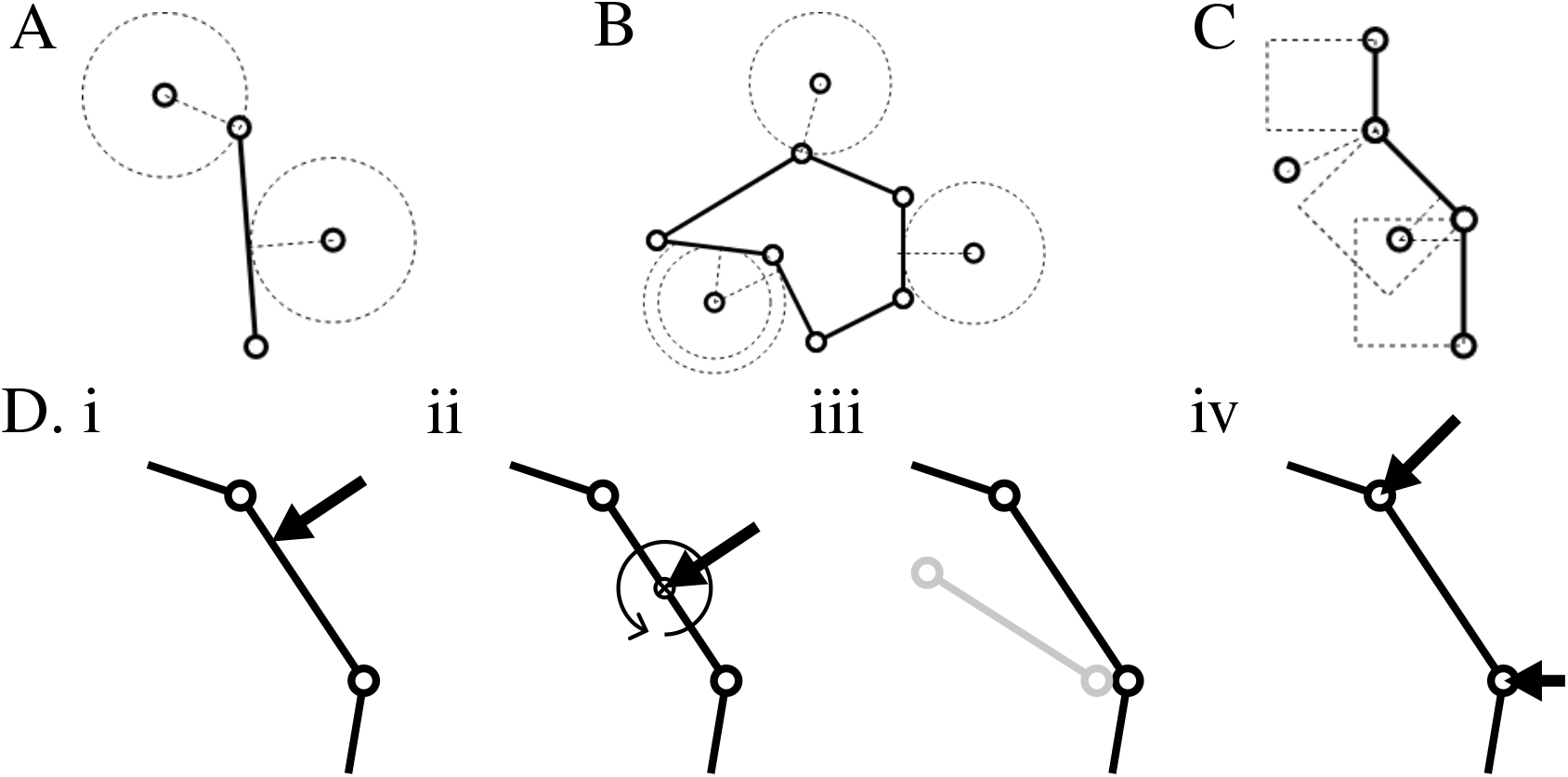
A schematic overview of the rigid body framework. A,B and C: Interactions between nodes and edges are determined by drawing a circle centred on the node and expanding its radius until it first contacts the edge. The place where it contacts determines the type of interaction. If a node is in proximity to a chain of connected edges (B and C), it can interact with multiple edges. But if it falls into the wedge shaped gap between interaction boxes (C), it will only interact with the shared endpoint. D: The process for resolving rigid body motion used in the models. Any force applied to the edge can be replaced with a force at the centre of drag and a moment (ii). This is then used to calculate the rigid body displacement that an isolated edge would experience in a given time step (iii). From this, we work backwards to determine an equivalent force that would produce the same resultant displacement of the nodes when they are treated independently of the edge (iv).

Interaction forces act along the vector pointing from the node to the contact point, while the force magnitude is calculated by using the length of this vector. Equal and opposite forces are applied to the node and the contact point on the edge. In a node-node interaction, the resultant forces are applied directly to the nodes. However, if the contact point is an interior point we must invoke rigid body mechanics.

Since we are assuming inertia is negligible compared to viscosity, we need to use a form of rigid body mechanics constructed around viscous drag coefficients. This is not something readily found in relevant texts, hence for the purpose of this work it has been developed from first principles. We introduce the important concepts in *Methods* and provide derivations in *Supplementary Material*. Figure 1D illustrates how rigid body motion is realised. Any force applied to a rigid body can be replaced with a force applied at the *centre of drag* (analogous to the centre of mass), and a moment (Figure 1D ii). We use the viscous rigid body equations of motion to determine the displacement of the edge in a given time step (Figure 1D iii), and then extract two equivalent forces that result in the same displacement through purely linear motion of the endpoints (Figure 1D iv). The equivalent forces due to edges, as well as any direct or internal forces are then summed for each node, and the simulation is evolved in time using the usual equation of motion (Equation (1)).

To illustrate the capabilities of our rigid body framework, we present three application exemplars: a model using isolated edges to represent bacteria, a polygonal cell model used to simulate a tumour spheroid, and an augmented vertex model applied to epithelial buckling where the layer can self-interact.

### Bacteria Exemplar: Mechanical interaction of cells can cause ordering of cell populations

At the simplest level, we can use the rigid body framework to produce an “over-lapping rods” model, using isolated edges to represent cells like bacteria and yeast. The inherent length of these cells means that a node-based framework is not suitable. Existing models have worked around this problem using phenomenological movement procedures [28] or by applying inertial equations of motion [32] to rod-shaped cells. The model by Smith et al. [30] uses a process that moves overlapping cells using an iterative procedure to minimise the total amount of overlap in the population. Simulations can advance rapidly depending on the hardware used, however, there is no guarantee of convergence, meaning the simulation may have unrealistic levels of overlap. Conversely, models that use inertial equations of motion may be less appropriate considering the systems are usually viscosity-dominated.

In 2008 Volfson et al [32] examined cell behaviour in a confined environment by constraining *E. coli* cells to a monolayer in a microfluidic channel. They were able to demonstrate that the spatial constraints of the environment influenced the orientation of the cells; specifically, cells tended to align with the axis of the channel. Here, we can replicate the experiment using the over-lapping rods model which is based on the viscous equations of motion. Cells are modelled as rods with a basic cell cycle that controls their growth. Internal forces drive the cell to target length via a spring force based on the length of the rod. Division replaces a cell with two daughter cells covering the same space (Figure 2D) and interactions are determined via neighbourhood searching (Figure 2E). Contact inhibition is implemented to prevent growth commencing when a cell is over compressed. Full details are given in *Methods*. We run 20 simulations each with a random seed and calculate average values.

**Figure 2:**
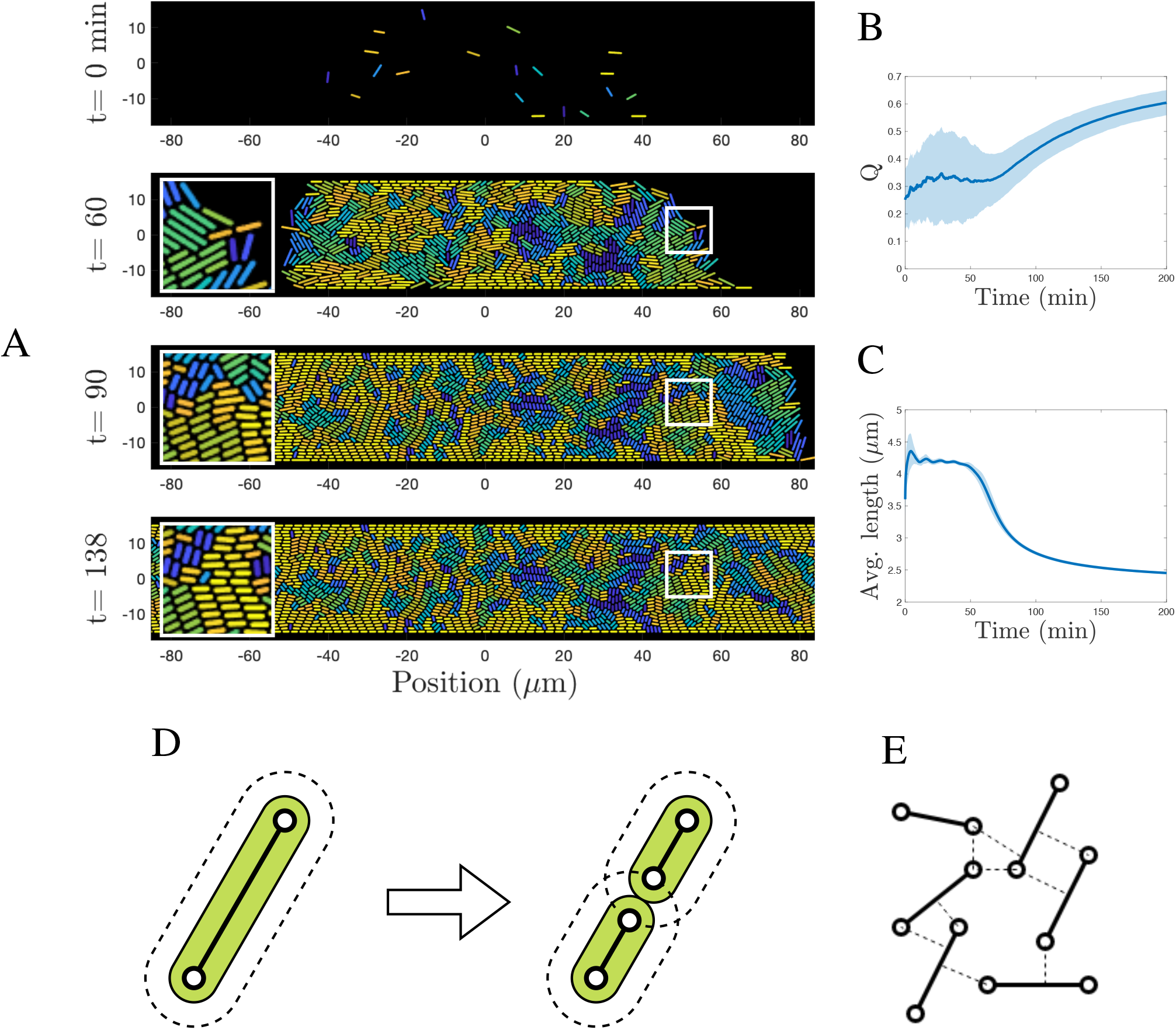
A simulation replicating the experiment by Volfson et al. [32] with microbes proliferating in a monolayer, constrained to a channel. A: Stages of the simulation showing times *t* = 0, 60, 90 and 138 minutes. Colours indicate the orientation of the cell (yellow is horizontal, blue is vertical). Insets show cells aligning with the channel axis. A video of the simulation can be found in the *Supplementary Material*. B: The order parameter *Q* that uses the orientation of the cells to quantify the disorderliness of the microbe population. The blue line is the average over 20 simulations, and the shadow represents one standard deviation. C: The average length of all cells in the channel. For small populations where the cells are loosely contacting, the average is near to the average of new and fully grown cells. As the channel fills up, the over-crowding restricts their ability to grow and the average length decreases. D: The division process for the over-lapping rods model. The extent of the cell is coloured in green. The dashed line shows where an adjacent node or edge will sit at equilibrium. E: A collection of isolated edges illustrating the set of interactions for this configuration.

#### Simulations and results

Using this model, we are able to reproduce several features observed by Volfson et al. [32]. Figure 2A shows several snapshots of our model. Initially, cells are randomly placed in the channel, with a random orientation. As the population grows, the cells increasingly make contact with the boundaries, causing them to align with the channel axis. This alignment propagates into the channel interior, particularly at the proliferating front.

In Volfson et. al. the authors use the order parameter

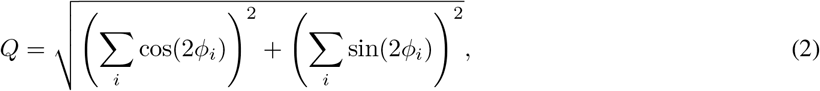

borrowed from liquid crystal theory [8] to measure the orderliness of the cell population. Here *ϕ*_*i*_ is the angle cell i takes from horizontal. This captures whether the cells are oriented in the same direction (*Q* ~ 1), or have a random orientation (*Q* ~ 0). Figure 2B shows how *Q* changes with time, averaged over 20 simulations, each starting with a population of 20 randomly oriented cells. During the initial stages, the orderliness varies quite widely, but on average is around *Q* ~ 0.3. From approximately *t* = 60 minutes, the orienting influence of the walls starts to propagate, leading to a higher overall orderliness. This qualitatively agrees with the findings by Volfson et al. The authors also note that as time progresses, the average size of the cells decreases. Figure 2C shows this behaviour is replicated in our model. This is likely a result of contact inhibition preventing growth from starting, and increased compression due to the densely packed cells.

The behaviour we see here is not unique to the over-lapping rods model. However, this is the first model to directly apply viscous rigid body mechanics to the motion of bacterial cells. It may offer more physical realism compared to other approaches, which could have implications for other emergent population level behaviours.

### Spheroid Exemplar: Cell growth and movement is slowed by compression from surrounding tissue

Using the rigid body framework, we can build a cell model that combines the the clear definition of cell area and shape of the vertex model with the freedom of movement seen in the over-lapping spheres model. We construct a cell using a collection edges to form its boundary, but rather than sharing these with adjacent cells (as is done in the vertex model), each cell has its own set. Inter-cellular forces use the node-edge interaction mechanism, allowing smooth reactions and deformations, while also permitting total separation of cells.

A suitable pedagogical setup to test this model is a tumour spheroid. Spheroids are among the simplest biological systems to translate into a computer model. They have been examined in many works, from a mechanical investigation of growth with oxygen diffusion [5], to detailed models that include vascularisation [1]. In the wet lab, tumour spheroids are used to investigate the early behaviour of cancer [13]. A cell line is allowed to grow *in vitro*, creating a cluster of cells that consume nutrients and oxygen as they proliferate. A common observation is the halting of proliferation in the core of the spheroid as its size increases, which is understood to be caused by dwindling resources and increasingly dense packing of the cells [13].

Here, we model the growth of a tumour spheroid using decagon cells built from edges. Interactions between cells are handled by the node-edge interaction mechanism (Figure 3F), while internal forces are handled using energy methods akin to those used by Nagai and Honda [20]. Cell growth is handled by a basic cell cycle model, and division is modelled by cutting the boundary into two halves and turning the halves into new cells by adding edges (see Figure 3E). We also implement contact inhibition to prevent cells from commencing growth when they are under high compressive stresses. Full details are found in *Methods*. We run 20 simulations from random seeds to calculate average values.

**Figure 3:**
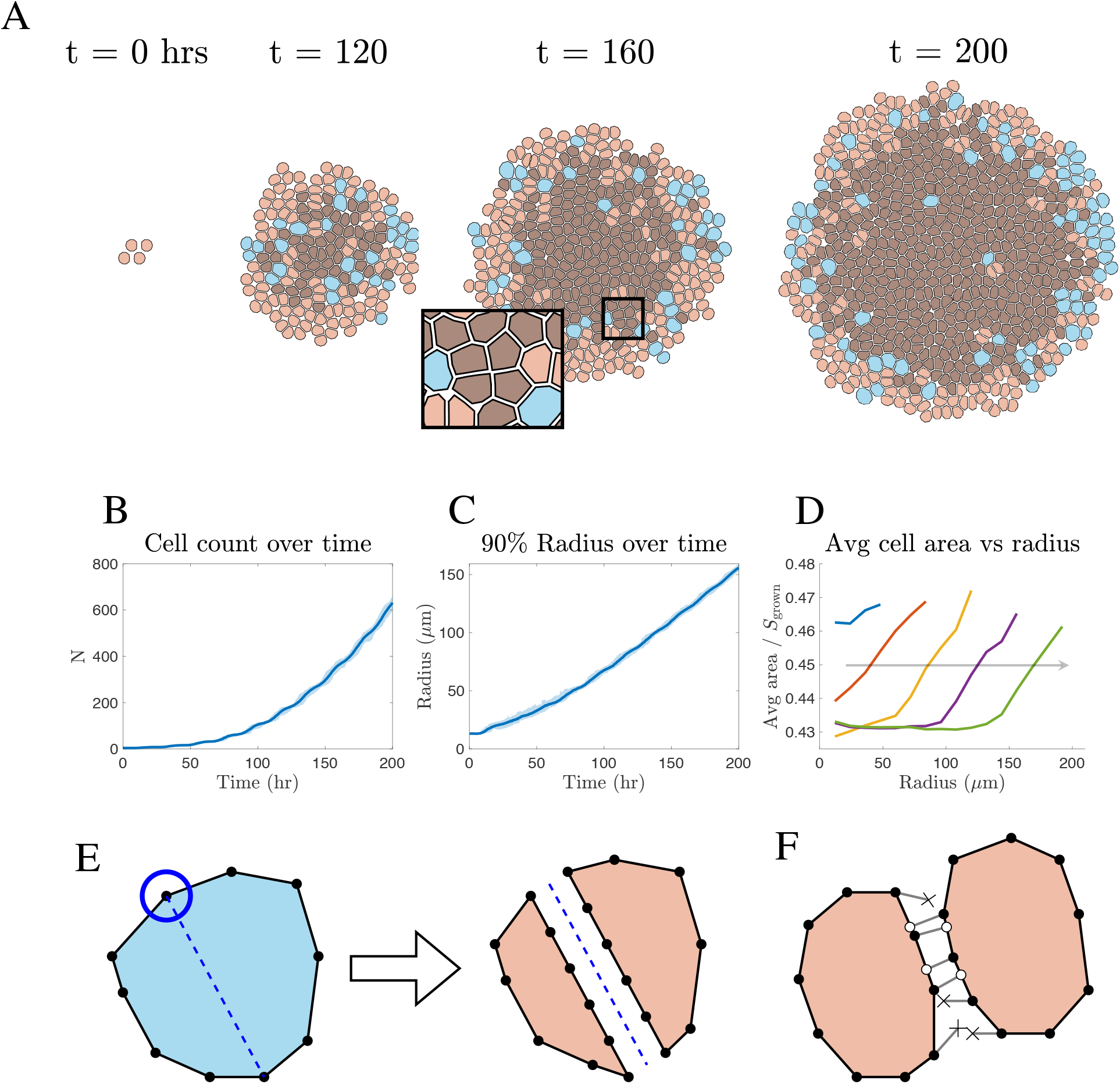
A tumour spheroid simulation using the rigid body framework to represent each cell as a decagon. A: Time snapshots of the growing spheroid. Pink cells are in a non-growing resting state after division, while blue cells are actively growing. As the spheroid grows, the compressive forces from the increasing cell mass prevent the core cells from commencing their growth phase (dark coloured cells). Inset: A spontaneously forming rosette. A video of the simulation can be found in the *Supplementary Material*. The plots show statistics averaged over 20 simulations for B: Cell count over time, C: the radius capturing 90% of the cell centres over time, and D: the average cell area as a fraction of *S*_grown_ versus radius from the centroid at times *t* = 40, 80, 120, 160 and 200 hours. Grey line is the contact inhibition threshold stopping growth. Arrow indicates increasing time. E: The division process. A random vertex is chosen and the cell is split to evenly share the edges between the new cells. Additional edges and nodes are added to keep the edge count constant for all cells. The new nodes and edges are placed so the cells start at their preferred separation. In the subsequent time steps, the shape of the cells quickly adjusts to be roughly similar F: A schematic showing how interactions between neighbouring cells are determined. Open circles represent contact points, and crosses represent potential interactions that are too distant.

#### Simulations and results

Figure 3A shows the time progression of a spheroid using this decagon cell model. The snapshots show that as the tumour grows in volume, the interior cells become too compressed to start growing, while the cells in a band around the perimeter are able to grow and proliferate.

Figure 3B shows the growth in cell number, while Figure 3C plots the radius capturing 90% of the cell centres, both averaged over 20 simulations. After the cell population has reached a reasonably circular shape (roughly *t* > 100 hours), the radius grows at an approximately constant rate, implying the cell population is increasing quadratically.

Figure 3D shows the average cell area as a function of distance from the spheroid centre. As time progresses, the inner compressed core expands, while proliferation is limited to a narrow band around the rim. These population level behaviours agree with other models and experiments [5], [13]. We also see some features here that may not be possible in other models. The inset for *t* = 160 hours of Figure 3A shows a spontaneously forming rosette, where four or more cells form a junction. In a vertex model, rosette appearance needs to be specifically included in the model by modifying the way node swapping is handled, whereas here it has formed as a natural consequence of the interaction mechanism. More generally, we see cells are able to take a range of shapes depending on their local environment. Cells closer to the tumour growth front are less compressed, hence are closer to a regular decagon in shape. Cells closer to the middle have a more varied shapes, some having an elevated aspect ratio, others approaching rectangles or pentagons.

### Epithelial Exemplar: Smooth self-interaction of tissues can capture large scale deformations

One of the most exciting capabilities of the rigid body framework, is its ability to allow tissues to interact without the need for interaction procedures that limit the scale and realism of the model. As an example we look at a deforming epithelial monolayer.

Epithelial monolayers are a common tissue found in the human body and throughout the animal kingdom [10]. Due to their simplicity, monolayers have attracted much attention from cell-based modellers, with common questions relating to the buckling behaviour of the layer due to cell growth. Many models allow an epithelial layer to grow and buckle ([22], [12] and [15] are but a few), but the buckling can usually only continue as long as no self-interaction occurs. Very few models have been able to produce simulations that can meaningfully continue for long-term large-scale self-contacting deformations. This has meant that modelling morphogenesis beyond self-contact has been difficult to achieve. Merzouki et al. [19] use a phenomenological procedure in their epithelial vertex model to handle self-contact. When a node (vertex) moves into an adjacent cell, the nonphysical situation is resolved by moving the node back to the edge that it crossed. This prevents unrealistic cell overlapping, however, the algorithmic response means that forces are not transferred between interacting cells. Such behaviour causes energy to be lost from the simulation that might otherwise have produced appreciable movements. A mechanical resolution to the interactions is therefore more desirable.

Here, we implement an epithelial monolayer vertex model similar to that used by Merzouki et al. Rectangular cells are joined in a contiguous ring, and follow the basic cell cycle model described in *Methods* to control their growth and division (Figure 4C) with the same parameters as the polygon cell model. Most importantly, we use the node-edge interaction mechanism to model interactions when the monolayer self-contacts (Figure 4D).

**Figure 4:**
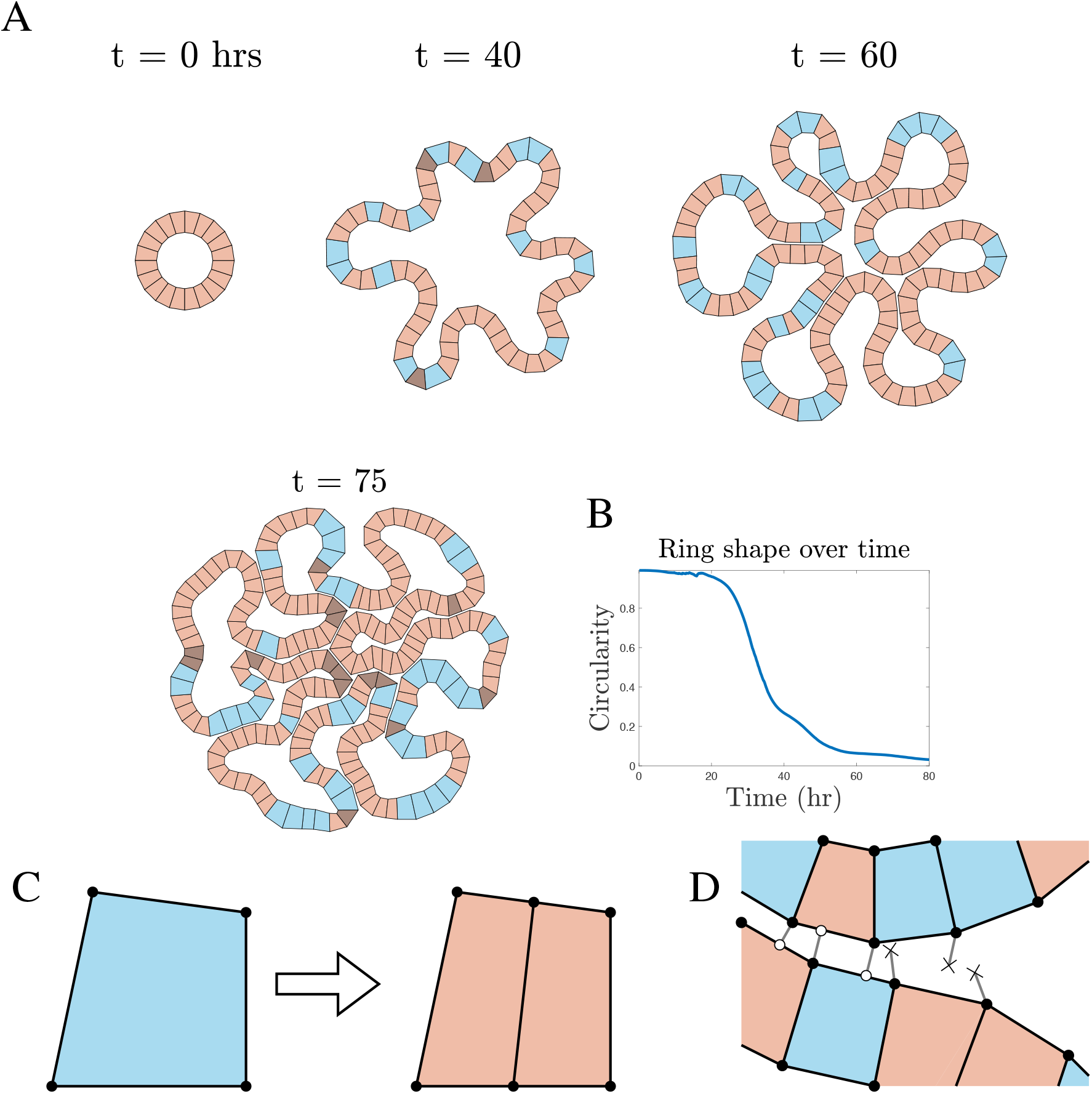
A simulation of a buckling ring of epithelial cells. Pink cells are in a non-growing resting state, blue cells are actively growing, and dark coloured cells are contact inhibited. A: Several time snapshots of the model showing initial buckling, and increasing levels of self-contact. By *t* = 75 the epithelial layer has progressed to a point where it has self-contacted in numerous areas. The node-edge interaction mechanism allows the cells to transmit forces at these contact points and slide past each other freely. A video of the simulation can be found in the *Supplementary Material*. B: The circularity measure of the ring of cells. The simulation can run well past initial contact, to the point where circularity approaches zero. C: The simple division process for a fully grown cell. When the cell is ready to divide the top and bottom edges are split exactly in half. An edge joins the new nodes together, thus creating two cells. D: A schematic illustrating how the interactions between the layer are found. Open circles indicate contact points, and crosses show potential interactions that are too distant.

#### Simulations and results

Figure 4A shows the progression of this model as it grows and buckles. Early times show the usual behaviour, where the layer buckles producing finger-like extrusions. Around *t* = 60 hours, we start to see the first stages of self-contact, where the layer is interacting with itself, but the deformation is small. By *t* = 75 hours (Figure 4A) the node-edge interaction mechanism has allowed the layer to become extremely warped and still progress in a mechanically consistent way. The interactions are able to dynamically reconfigure themselves as nodes slide past, without causing them (or the edges) to become stuck, all while allowing movement energy to be transferred across the contacts.

To track the state of the simulation we calculate the circularity of the monolayer

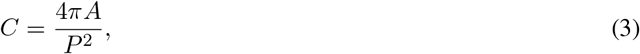

where 0 ≤ *C* ≤ 1 and *C* =1 for a circle. Here *A* is the internal area of the ring, and *P* its internal perimeter. We see that the node-edge interaction mechanism allows the simulation to progress to the point where circularity approaches zero (Figure 4B).

Other models have relied on identifying collisions and resolving them using a procedure to handle self-contact. This can be computationally intensive if implemented naïvely, although efficient algorithms do exist. Critically though, these procedures do not account for force transfer between components, limiting the realism of the model. By introducing the node-edge interaction mechanism, forces can be transferred using physical principles, leading to more realistic motion of contacting tissues. Additionally, since the node-edge interaction mechanism functions regardless of cell connectivity, the epithelial layer can interact with other objects in the environment, such as supporting structures. This opens up the possibility of modelling how epithelial layers interact with their underlying stromal tissue to produce dynamically stable structures, something that has been identified as an underdeveloped area in cell based modelling [2].

## Discussion

### Capabilities

In this work we have presented a new modelling framework that expands the possibilities for cell-based modelling by introducing a mechanical way for edges to interact with their environment in a system dominated by viscous drag forces. It can be seen as the natural progression from node based modelling, where the foundational object is now a one-dimensional edge, rather than a zero-dimensional node. This is significant because edges offer a much more intuitive way to model objects with length compared to a node-based framework.

The first model presented was perhaps the simplest application of the rigid body framework, namely the over-lapping rods model. It produced a way to model rod shaped cells where interactions are based on the viscous equations of motion, rather than phenomenological procedures. This is something that has not been available to researchers up until now. Existing approaches have drawbacks in computational complexity (i.e. searching for collisions, and iterative solutions), and mechanical realism (i.e. forces are not transferred consistently, or are based on inertial motion). The over-lapping rods model successfully addresses these.

We are also able to join edges together to produce contiguous surfaces. Existing models construct surfaces out of nodes. In this situation, there is fundamentally nothing stopping one node from moving between any pair of adjacent nodes. This is a major problem for modelling barriers since their function is to separate one region of space from another. There are ways to address this problem within a node-based framework, one obvious way being to increase the density of nodes on the surface, however this clearly comes at a computational cost, and it may not necessarily behave as intended. If a node approaches a barrier built from edges, the barrier can react as if a force is applied directly to its interior, rather than approximating the interaction directly with the nodes. As a consequence, edges can cover suitably large regions of space and still provide a barrier response.

Another benefit of the rigid body framework is that we can model cells with arbitrary shapes without needing to consider node sharing. In a vertex model, cell shape is clearly defined, however, the need to share nodes between multiple cells requires special processes for cells to move through a tissue. As a consequence not all common types of cell motion are possible, in particular, there is no simple way for cells to migrate along surfaces. With the polygon model, cells can easily move through tissues, without needing to consider how nodes are shared, and can even become completely isolated from other cells while maintaining their shape. All interactions are determined by neighbourhood searching, meaning surface interaction is built into the model. This means barriers, such as membranes, or tissue surfaces can easily be implemented, with minimal extra effort. This has been demonstrated in the epithelial monolayer model where a simple vertex model was augmented with the node-edge interaction mechanism to allow the tissue to self-interact. It is a relatively small step to extended this to include separate interacting tissues that are internally governed by a full vertex model. This has important applications for modelling morphogenesis (for example, neurulation [29]), and for modelling tissues where an epithelial layer may be tightly folded (such as the colonic crypt [14]).

Neighbourhood searching also has benefits in terms of computational time. Efficient and relatively simple algorithms exist for finding nearest neighbours that scale linearly with number of nodes and number of neighbours (i.e. *O*(*kn*) time) [3]. It can be shown (see *Supplementary Material*) that edges do not appear to add any extra computational burden to neighbour searching, and in fact they may provide an advantage in certain scenarios.

As with all multi-cellular modelling, the rigid body framework is not without its own drawbacks. The over-lapping rods model suffers from the same poor definition of shape as the over-lapping spheres model, especially for highly confluent layers. It also has the risk of rods crossing, creating a non-physical situation, although this is mitigated by choice of force law. For polygon cells, special attention is needed to ensure the boundary remains a simple polygon, where it does not cross itself, and this can be achieved by implementing additional forces to oppose such motion.

### Future directions

In the models presented here, we have explored only a handful of the potential applications for a rigid body framework. The tumour spheroid and epithelial monolayer models show how edges can be used to robustly model cell surfaces, however there is another major biological feature that can exploit this behaviour, namely, membranes. Membranes are an important part of organs that receive a lot of modelling attention (for instance the intestinal crypt [12]). Despite this, the role of the membrane is often neglected by assuming it provides an immutable surface [31], [11]. This is a sensible simplification when the behaviour of interest does not rely on the dynamic shape of the crypt. But if the behaviour is fundamentally tied to deformation (i.e. crypt fission, or polyp formation), then a membrane model that can handle large scale movements is vital. There are currently few models that attempt to fill this gap in two dimensions, and even fewer in three dimensions; the rigid body framework is ideally positioned to address this. Smaller scale biological systems may also be amenable to the rigid body framework, for instance the cyctoskeleton made up of long slender microtubule and actin filaments [25], or even the small scale behaviour of basement membranes and basal laminar, made up of a collagen fibre network [26].

Perhaps the most promising future application is in three-dimensional multi-cellular modelling. Large scale three-dimensional modelling, with many thousands of cells, is well served by the over-lapping spheres model. Smaller scale modelling where the structure of a single cell is the focus is well served by sub-cellular and finite element methods. The intermediate scale, where cell shape can have important consequences, but where the number of cells makes the more detailed approaches intractable, is currently underdeveloped. Vertex models and tessellation models have been adapted to three dimensions for modelling dynamic tissues [23], or reconstructing tissues from images [6], but as with their two-dimensional counterparts they are fundamentally built upon nodes. This leaves them ill-suited to constructing barrier objects. While as yet undeveloped, there is a clear path to adapt the rigid body framework into three dimensions by expanding viscous rigid body mechanics to two-dimensional surfaces.

The major strength of this development is that it is not a model in and of itself, rather it is a building block from which a new branch of multi-cellular modelling approaches can emerge. Existing models can be integrated into the rigid body framework to expand their capabilities, as seen for the vertex model. It can also be used as the basis for a new model in its own right, as seen with the over-lapping rods model. There is a wide range of possible further applications that can exploit interacting edges as well as many avenues for building on and improving the models presented here.

## Methods

### Application details

#### Bacteria model

For this application, each cell is represented by an isolated edge, where rotation is captured by equivalent forces (Figure 1D iv), and interactions between two endpoints are handled like independent node-node interactions. We use a standard force equation to calculate the magnitude of cell-cell interaction forces (See Equation (21)). In this model we only consider repulsive forces between the cells by setting the attraction force parameter, *s*_a_ = 0. According to Volfson et al. an *E. coli* cell can have a length from 2 to 5 times the cell diameter (CD where 1CD ~ 1*μ*m) depending on the point in its growth cycle. [32]. Cells will experience an internal spring force that will push them to a given target length according to Equation (28). During the course of the simulation, cells will not often reach their target size, hence to achieve the required range of aspect ratios and therefore observe the ordering behaviour, we must tune the target length. Here it is necessary to set a fully grown cell’s total target length to 6CD which is made up of *S*_grown_ = 5CD for the length of the rod and 1CD from the preferred separation distance around the ends of the cell. A newly divided cell will have a target size of 3CD and therefore a target rod length of *S*_0_ = 2CD. Average cell length is plotted in Figure 2C demonstrating this choice achieves the required aspect ratio range.

We apply a standardised cell cycle model designed to control the target size of the cell (see *Cell Growth and Division* below). This is different to those used in other works for rod-shaped cells, but we use it here for consistency with the other applications. A new cell starts off in a resting state where the target rod length is constant at *S*_0_. It remains in this resting state until it reaches the age *t*_0_ which is chosen from the uniform distribution *t*_0_ ~ *U*[0, 2] minutes. If the measured size of the edge is at least *s* ≥ *γS*_0_, where *γ* = 0.9, then the cell is allowed to start growing, otherwise it waits in a contact inhibited state until the size requirement is met. Growth is modelled by linearly increasing the target size from *S*_0_ to *S*_grown_ over a period of time *t*_g_, which is also chosen from a uniform distribution so that *t*_g_ ~ *U*[8, 12] minutes. Division occurs immediately after *t*_g_.

Division produces two collinear, equal length daughter cells that fill the same space as the parent cell. This is achieved by splitting the edge into two new edges with a small gap separating them at the preferred separation distance (Figure 2D). Cells arranged like this may remain perfectly collinear in the absence of outside forces, but we prevent this with addition of random noise when the motion is calculated (see *Practical model implementation details*). After division, the cell cycle for both cells starts from *t* = 0, hence the new target size will be *S*_0_.

To run the simulation, cells are placed on a plane, with a top and bottom boundary making a channel 30CD in width, matching the 30*μ*m channel of Volfson et al. [32]. The boundaries are implemented by a simulation “modifier” that shifts endpoints vertically to sit exactly on the boundary if they happen to cross it. The left and right of the plane are open, and cells are free to travel in these directions without restriction. The simulation is seeded with a sparse collection of 20 randomly oriented cells placed within the central 80CD of the channel, with the cells starting at random point in their cycle. The parameters from Table 1 are applied and the simulation is allowed to run to *t* = 200 minutes. We run 20 simulations, each from a different random seed, and use them to calculate average population behaviours.

**Table 1:**
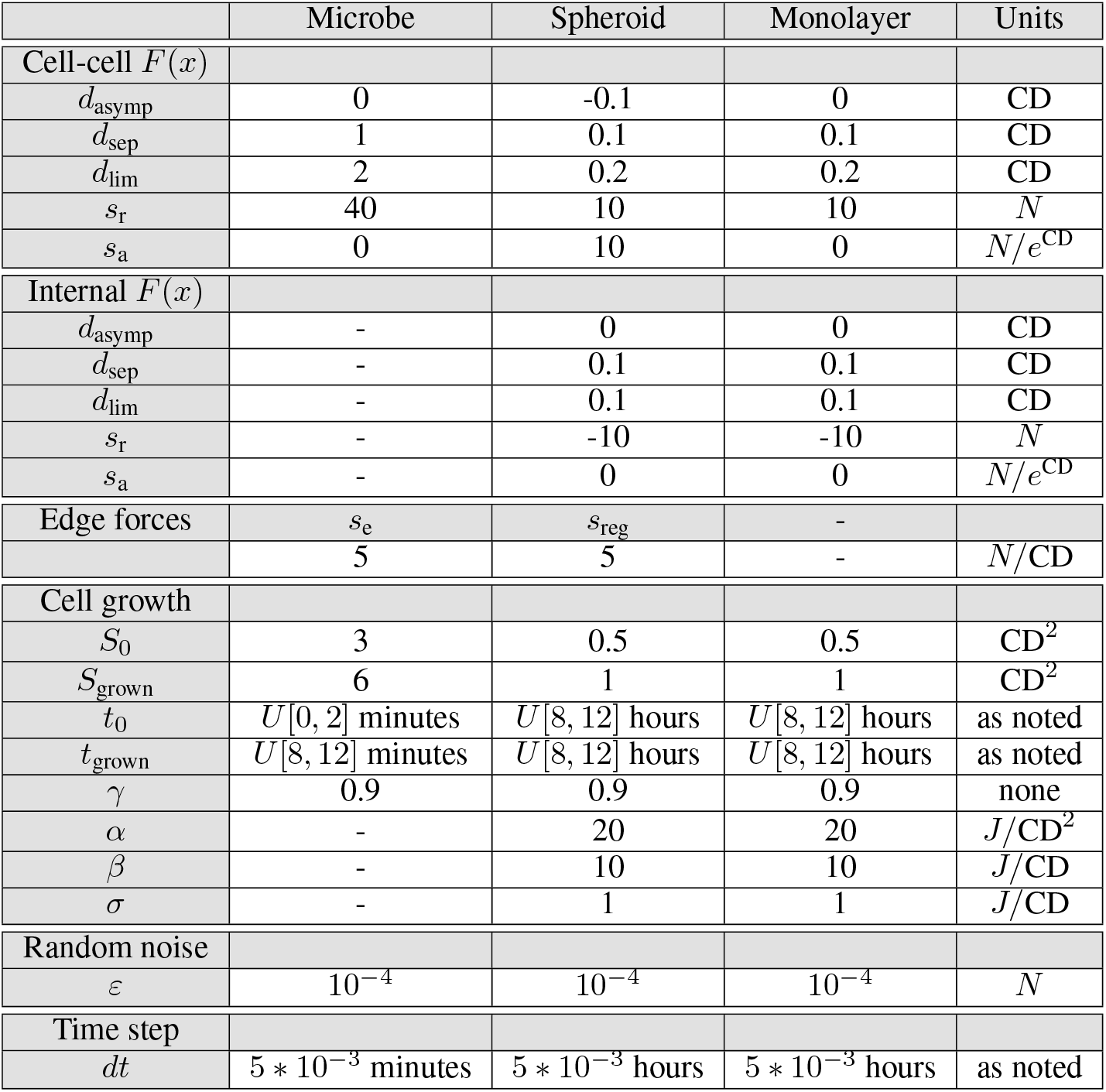
Parameters for each model described in *Methods*

#### Tumour spheroid model

Using the rigid body framework, we investigate a tumour spheroid by using an arbitrary decagon to represent a cell. The boundary of the cell is made up of a contiguous ring of edges, hence each edge shares its endpoints with two others. Interactions between cells (Figure 3F) are determined using the node-edge interaction mechanism. For computational efficiency, direct interactions between nodes are not considered, and this does not affect the results as discussed in *Practical model implementation details*. Forces between nodes and edges are calculated using a standard force law (Equation (21)), and for illustrative purposes, we allow cells to over-lap by setting *d*_asymp_ < 0. Cells are driven to a target shape using the the energy methods from Equation (29). The target cell area is controlled directly by the cell cycle model, and the target perimeter is driven by the target area by assuming the cell wants to form a regular decagon. The parameters used are collected in Table 1.

In this model a cell maintains a constant number of edges around its boundary during its life cycle. It is possible to make the number of edges depend on cell size for the sake of computational speed or shape accuracy, however this requires special attention to how cell boundaries are modified, and is left for future work.

We use the standard growth focused cell cycle model to control the target area of the cell. A new cell starts off at with target size *S*_0_, and remains at this target size until *t*_0_, which is chosen from a uniform distribution such that *t*_0_ ~ *U*[8, 12] hours. Contact inhibition holds the cell in this resting state until *s* ≥ *γS*_0_, where in this case *γ* = 0.9. Growth occurs over a period of time *t*_*g*_ ~ *U*[8, 12] where the target area increases linearly from *S*_0_ to *S*_grown_. Division is allowed to occur immediately after growth is complete, producing two daughter cells occupying the same space as the parent cell. We choose a cell diameter CD= 15μm to roughly match the size of a standard HeLa cell[4]. New cells will have a target size of *S*_0_ = 0.5CD^2^ and fully grown cells will be *S*_grown_ = 1CD.

For a polygonal cell, the process of creating new cells through division has several steps (Figure 3E). A random vertex is chosen to define the division axis, and the cell is split in half so that each daughter cell takes five of the edges from the parent cell. Five new edges of equal length are added in a straight line between the loose ends of each half to close the loop and form the new cells. The newly divided daughter cells will have an area slightly different from half of the parent cell’s area and a perimeter somewhat longer than the target perimeter due to the offsetting. This causes the new cells to reconfigure themselves over several subsequent time steps into a form that more closely resembles the physical shapes observed *in vitro* post division.

By using this simplified method of cell division, the distribution of edges around the boundary of the cell can have a significant impact on the shapes of the newly divided cells. In a worst case scenario, this can leave one cell with most of the parent cell’s area, and the other may be just a small sliver, potentially causing numerical issues due to unbalanced forces. However, this is substantially mitigated by applying a regularising force that drives the edges to be of roughly equal length while the cell is growing. Additionally, any disparity between daughter cells upon division is quickly lost due to the internal cell forces.

To initialise the simulation, four decagon cells are placed next to each other on a plane starting with their internal clock set to a random time between 0 and *t*_0_. The parameters from Table 1 are applied, and the simulation is allowed to run to *t* = 200 hours. We run 20 individual instances, each starting from a unique random seed, and use them to calculate average population behaviours.

#### Epithelial monolayer model

To examine a buckling epithelial monolayer, we integrate a simple vertex model similar to that used by Merzouki et al. [19] into the rigid body framework. Here, a monolayer is represented as a contiguous ring of quadrilateral cells. Each cell is made up of four edges, with the inner and outer edges being exposed to the external environment, and the clockwise and anticlockwise edges being shared with the adjacent cells. The cell behaviour is much the same as with the tumour spheroid model. The shape of the cell is controlled by embodied energies (Equation (29)), and the growth of the cell is controlled by the cell cycle with resting phase duration and growing phase duration both taken separately from the uniform distribution *U*[8, 12]. Interactions between adjacent connected cells are handled by applying internal cell forces to the shared nodes. Critically, interactions between adjacent non-connected cells are calculated using the node-edge interaction mechanism. Unlike the spheroid model, these interactions are repulsion only, so we set the attraction parameter *s*_a_ = 0. We will also not allow edges to overlap, hence we set *d*_asymp_ = 0.

When a cell in the monolayer has satisfied the conditions for division (the same as in the spheroid model), the cell divides by breaking both the top and bottom edges into two connected edges exactly half the length of the original edge. A new internal edge is added by joining the two new endpoints from the top and bottom (Figure 4C), thus separating the cell into two connected daughter cells of similar area.

To run the simulation, the monolayer is constructed by forming a closed ring of cells. The thickness of the ring is chosen to be 1CD, and the internal radius of the ring is set so that each cell starts with an area *S*_0_ = 0.5*S*_grown_ = 0.5CD^2^. The ring is initially comprised of 20 cells, each starting with an internal clock chosen from *U*[0, *t*_0_], and is simulated for a period of 80 hours. A complete list of parameters is found in Table 1.

### Implementation details

#### Viscous rigid body motion

At the core of this framework, we are assuming that the system is over-damped, where drag forces dominate. Surprisingly, equations of motion for over damped rigid body mechanics are difficult to find, despite (or perhaps because of) the abundance of information on rigid bodies in inertial systems. For the sake of completeness, a derivation of the concepts introduced below can be found in the section *Derivation of viscous rigid body mechanics* in the *Supplementary Material*.

As soon as an object has length, it is necessary to consider rotation. There are four key concepts for systems where inertia provides resistance to rotation: total mass, centre of mass, angular momentum and moment of inertia. However, these concepts are ill-defined when inertia is neglected. A rigid body theory balancing applied forces with drag needs equivalent concepts that use drag coefficients instead of mass.

The first of these concepts, we define as the *total drag*, **η**_*D*_. This is the drag coefficient integrated over the body

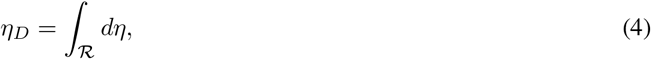

where 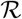 defines the body region. We also define the vector **r**_*D*_ that tracks the *centre of drag*, *D*, by

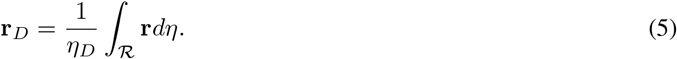

As with the centre of mass, the *centre of drag* is the point where we can consider the *total drag* of a rigid body (or system of particles) to be concentrated. If we apply a force that passes through the centre of drag of a rigid body, it is possible to show that there are no resulting rotations (See *Supplementary Material*), permitting us to treat the body as a point object. This allows us to describe the linear motion of a rigid body in an over damped system by the equation

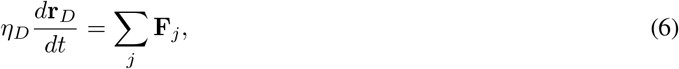

where **F**_*j*_ are the external forces applied to the body at *D*.

If a force does not pass through the centre of drag, then the body will also experience a rotation. When inertia is dominant, the rotation is resisted by the angular momentum, however when drag forces dominate, we use the quantity *angular drag*, *H*_*D*_, to resist motion, which in two dimensions is defined by the scalar

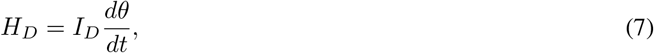

where **θ** is the angular position of the body relative to some axis, and *I*_*D*_ is its *moment of drag* about *D*. This can also be calculated in three dimensions, however we need to consider moments about all three axes, resulting in a vector form of *H*_*D*_. We leave this for future work. Continuing in 2 dimensions, we define *I*_*D*_ by

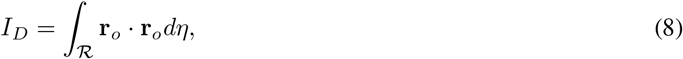

where **r**_*o*_ is the distance from *D* to the infinitesimal unit of drag *d*η**. The angular drag balances with the externally applied moments **M**_*k*_ giving us the second equation of motion

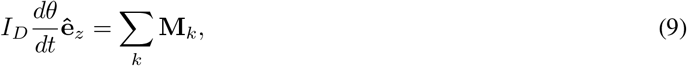

where 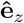 is a unit vector perpendicular to the plane of the cells. This allows us to determine the rotation of the body.

For a rigid body with a single externally applied force **F** that does not pass through through the centre of drag, it can be shown that **F** can be replaced by a force acting at the centre of drag and a moment [17]. Therefore, the motion of the body can be completely determined by evaluating the two equations

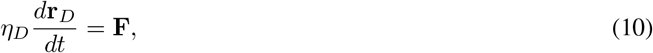

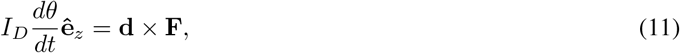

where **d** is a vector from the centre of drag to the line of action of **F** such that the two vectors are perpendicular.

#### Motion of an Edge

Figure 5 shows an isolated edge in an overdamped system subject to an external force. Vector **r**_*D*_ tracks the position of the centre of drag, and the vectors **r**_*D*1_ and **r**_*D*2_ track the positions of the rod’s endpoints relative to *D*. We can then construct vectors that track the position of the endpoints of the edge, **r**_1_ = **r**_*D*_ + **r**_*D*1_ and **r**_2_ = **r**_*D*_ + **r**_*D*2_. We only consider forces that act perpendicular to the edge, hence we introduce the force **F**_*A*_ which is applied to the edge at A, the point of action. To solve the motion of the edge, we need to carefully apply Equations (10) and (11).

**Figure 5:**
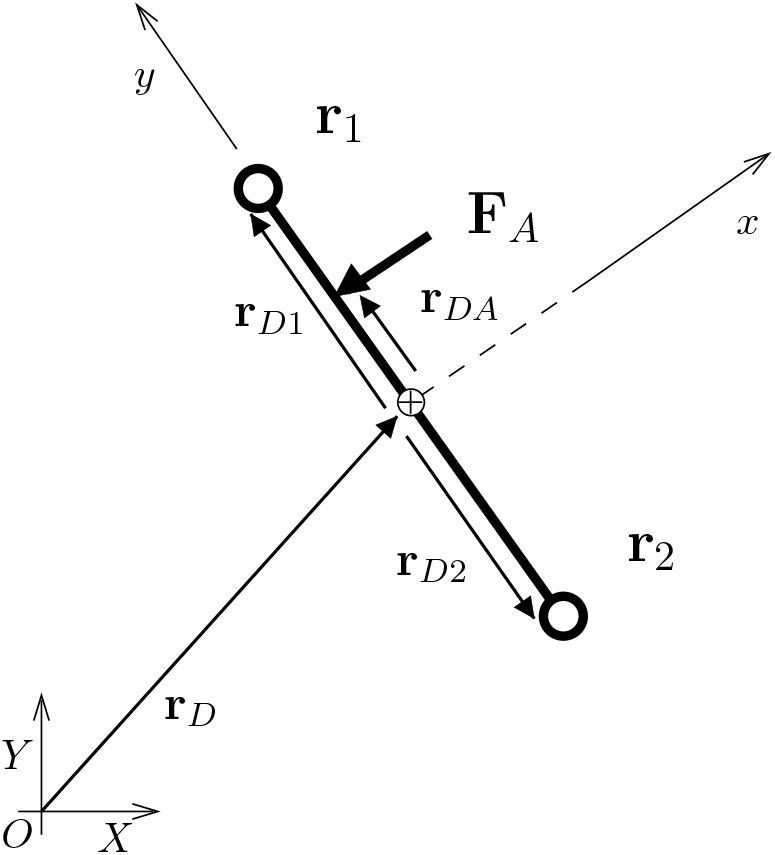
An isolated edge in an overdamped system with a centre of drag *D* (the crossed circle centroid symbol) and endpoints 1 and 2. A force **F**_*A*_ is applied at point *A*. The coordinate system (*X*, *Y*) is fixed in space, and the coordinate system (*x*, *y*) is rigidly attached to the edge. The *y* axis is fixed to the axis of the edge, and the *x* axis passes through the centre of drag. The edge (hence the axes (*x*, *y*)) rotate about *D* with angular velocity of magnitude *ω* pointing out of the page.

Given the vector quantities described above, we can expand out Equations (10) and (11) with the relevant scalar components. For the linear motion, we write the scalar components relative to the fixed system of coordinates (X, Y), and for the rotation it is convenient to write the components relative to the rotating system of coordinates (*x, y*), since we are interested in the position of the endpoints relative to *D*. Noting that *r*_*DA,x*_F_*A,y*_ = 0, these end up being

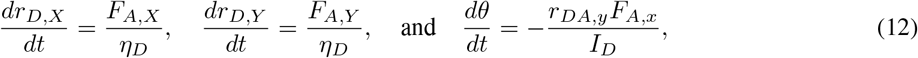

where the character following the comma in the subscripts represents the component in the respective coordinate system. The total drag, centre of drag, and moment of drag are calculated depending on how the edge is modelled. In the examples in this paper, we assume the drag is concentrated at the endpoints, hence

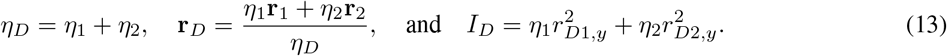

To actually solve these for the subsequent motion, we need to apply an intermediate step to deal with the rotation.

#### Converting moments into forces

As is commonly used when solving motion in cell-based models, we apply the Forward Euler method to determine the rotation and position at the next time step of a simulation [24]. The motion of the centre of drag is trivially solved at the next time step by

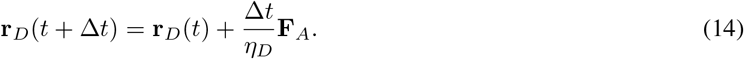

We can also quickly find the change in angle of the edge using this method,

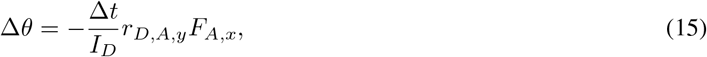

but to find the new position of the edge, we must use the rotational transformation matrix defined by

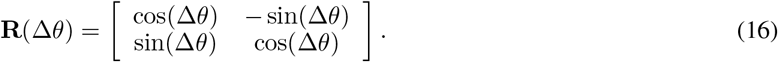

This rotates the body system of coordinates (i.e. the axes (*x*, *y*)) within the fixed coordinate system (i.e. (*X*, *Y*)). Since the position of the edge is uniquely determined by its endpoints, the goal of the procedure is to find the positions of the endpoints *j* at the next time step. Using the rotation matrix, the position vectors of the endpoints relative to *D* become

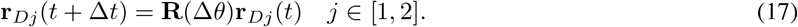

The final position of an endpoint due to the force applied to the edge is found by superimposing the two movements ^3^

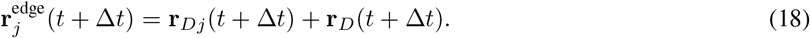

For an isolated edge with a single applied force, this is sufficient to solve its motion, so 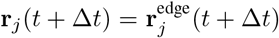 resulting in true rigid body behaviour. However, if the edge is connected to one or more other edges (as would be seen when modelling cell boundaries), rotation cannot be applied so easily. By moving the endpoint of one edge, we will necessarily be altering the state of any other edge sharing that endpoint. To accumulate the motion due to each edge, we define the *rod force*

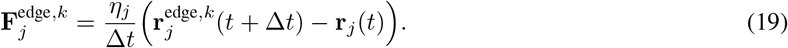

This is the equivalent force that would push the endpoint *j* of edge *k* to the position 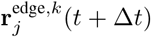 through purely linear motion. To collect the forces on an endpoint, we need to consider global indices that identify the given node and edge in the context of the whole simulation. If *J* represents any node, and *k* identifies each edge that has *J* as an endpoint, then we define the index function *j*(*J, k*), which identifies the position of *J* in *k* (i.e. whether it is endpoint 1 or 2). The total equivalent force on each endpoint, due to the rotation of the edges is then

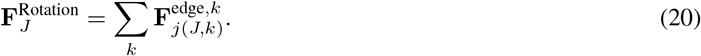

In effect, we have converted the rotation of an edge into a force that treats each endpoint as an individual point object. Under this definition, the edge can no longer be considered a true rigid body, since, depending on the forces applied, the edge can expand and contract in length. To describe this situation, we introduce the term *semi-rigid body*: the forces applied to the body come from treating it as rigid, but the resultant motion need not conserve the length of the edge. As a consequence, special care is needed during a simulation to ensure any given edge maintains a sensible length. Additionally, this makes integrating the whole system through time a simple task. We can use the same processes applied to node-based models since we have no requirement for directly handling rotations.

#### Finding interactions

A node-edge interaction occurs when the endpoint of one edge (a node) comes into proximity of another edge. In this situation, we can identify the closest part of the edge by taking a circle centred on the node, and expanding it until it first contacts the edge in question (Figure 1A and B). This point, which we call the *point of action*, will be used as the point where any interaction forces are applied. The node and the point of action taken together form the *line of action* for any subsequent forces. If the point of action is on the interior of the edge, the line of action will be perpendicular to the edge, and if the point of action is an endpoint, then the line of action extends radially out from the endpoint (see Figure 1A).

When the edges are connected and form an external convex feature as in the top of Figure 1B, the node will only interact with the endpoint. If on the other hand, the connected edges form a concave feature as in the left of Figure 1B, a node can interact with both edges, but will not interact with the endpoint. These situations can alternatively be viewed by drawing an interaction box around the edges as done in Figure 1C. The upper node is close to the two edges, but sits in a wedge where it can’t interact with the interiors, hence it interacts with the shared endpoint. The lower node is in two over lapping interaction boxes, hence it interacts with the interior of both edges and not the shared endpoint.

In either of these cases, two forces are applied along each line of action, one at the point of action, and the other at the node. They are equal in magnitude and opposite in direction.

#### Force magnitude

The magnitude of the forces is determined by the position of the node within the *region of interaction* around the edge. Figure 6 illustrates how the regions of interaction are formed around an isolated edge (Figure 6A), and an edge that forms part of a cell boundary (Figure 6B). The dashed line indicates a preferred separation locus. It is found at a constant distance from the edge, *d*_sep_. If a node is located on the locus, no force is applied. In effect, this defines the size and shape of the cell. Inside the dashed line is the repulsion region, where a force pushes the node and the edge apart. Between the dashed and dotted lines is the attraction region, where a force will pull a node and edge together. The dotted line defines the limit of interactions, and is found at a constant distance *d*_lim_ from the edge. Outside this limit, no interactions are calculated. The distance *d*_asymp_ represents the point where the force should asymptote to infinite repulsion. For an isolated edge (Figure 6C), we must have *d*_asymp_ ≥ 0. This prevents isolated edges from over-lapping, a situation which has no physical meaning. From a computational perspective, this is equally important for preventing a node getting pushed away on the wrong side of the edge since the direction of the force is dependent on which side the node approaches from.

**Figure 6:**
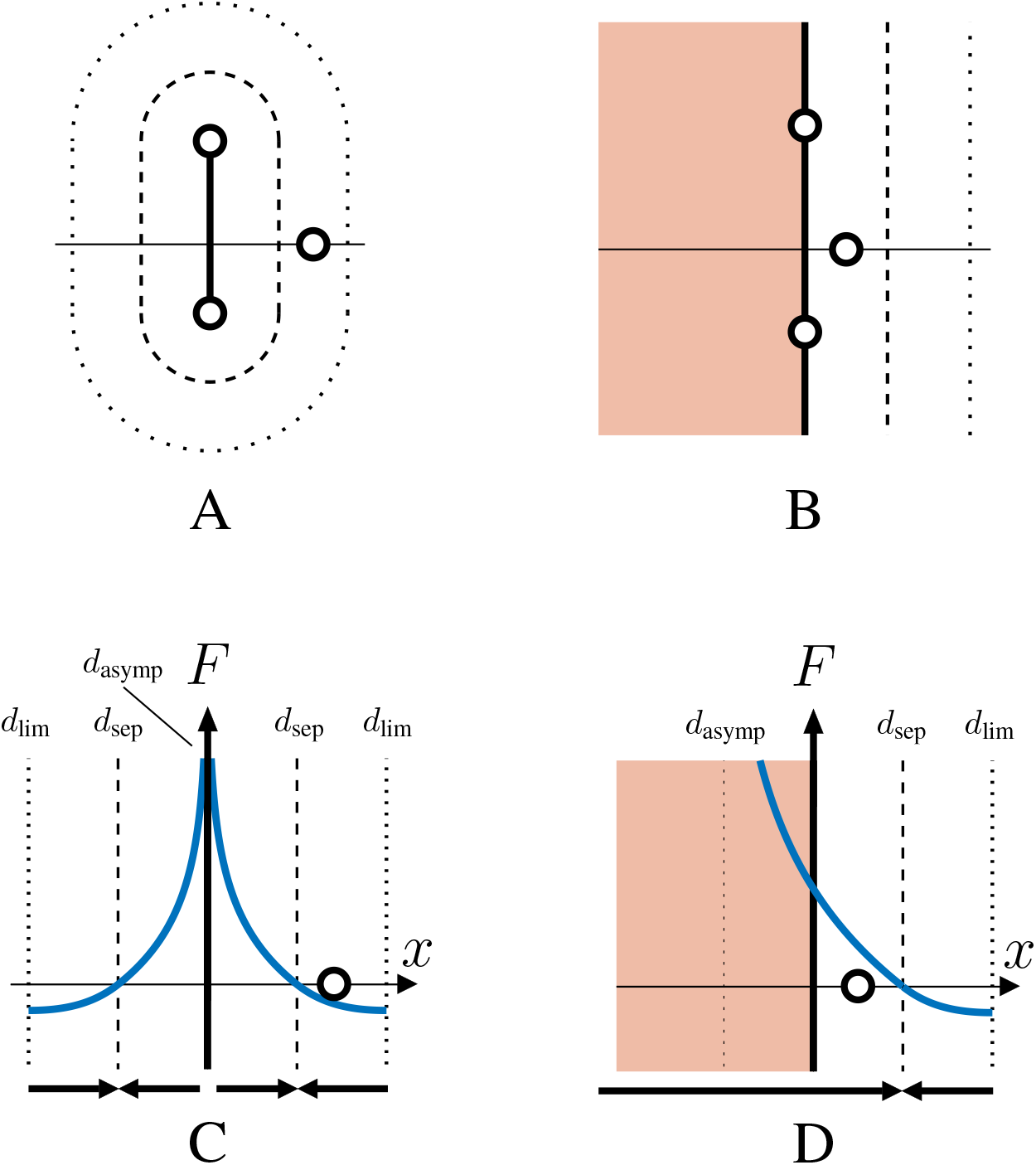
Interaction regions about an edge and illustrative force laws. A shows the regions around an isolated edge and B shows the regions around a cell with multiple joined edges. The thin horizontal line represents the line of action and is copied below in C and D as the *x* axis. The dashed lines represents the preferred separation locus (*d*_sep_), with nodes being pushed or pulled towards this line if they are interacting with the edge. The dotted lines define the limit of interaction (*d*_lim_). C and D show how the scalar value of the force is calculated as a function of perpendicular distance from an edge. Positive values represent repulsion, and negative attraction. In C, nodes cannot cross the edge (which coincides with the *F* axis) hence *d*_asymp_ = 0 In D, if a node is allowed to cross an edge, then the hard boundary will be found inside the cell, and is indicated by the internal dotted line (*d*_asymp_ < 0).

For an edge that is part of a cell boundary (Figure 6D), preventing over-lap is less important, since we have a clear way to orient the edge by using the inside of the cell. Depending on the specific force law, a node can be allowed to cross into the cell, or it can approach a hard barrier at the edge, hence we can choose *d*_asymp_ to have negative values. In either case, there is only one preferred separation point on the line of action, found outside the cell, so the direction of the force does not depend on which side of the edge the node appears.

A force law that encapsulates these requirements is

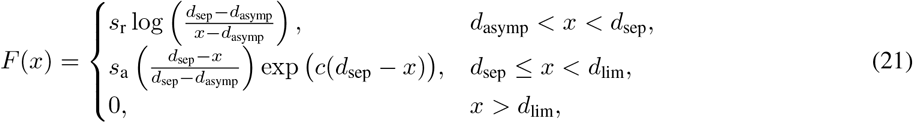

where *s*_r_ and *s*_a_ are parameters setting the strength of the force in the repulsion and attraction regions respectively. The parameter *c* controls the shape of the attraction force and in the models presented here will be *c* = 5. If *s*_r_ = *s*_a_, then the derivative of *F* is smooth at *d*_sep_. Other force laws can be used that capture the required features.

#### Applying a force to rods or boundaries

The force **F**_*A*_ was applied without specifying exactly where it came from. However, in the model we know that it comes from a node-edge interaction. In order to calculate the force, we need to consider the two possible cases, i.e. that the edge is isolated, or that it is part of a boundary.

If the edge is isolated and the force is due to being in proximity to node *J* (which is the endpoint of some other edge), then we will only consider |**r**_*AJ*_|, the length of the vector from *A* to *J*. This is due to the fact that we are not allowing isolated edges to cross. If we did allow edges to cross there is no simple way to determine how the forces should be applied in order to undo the crossing. We would need to track and record how each edge is oriented with respect to every other edge in its vicinity over multiple time steps. We will not attempt to do that here, so we only need to consider the magnitude of the vector separating the two points. The force is then determined by

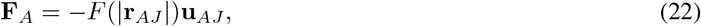

where *F* is the function defined in Equation (21), and **u**_*AJ*_ is the unit vector from the point of action *A* to node *J*, which is normal to the edge. The negative sign is necessary because we take repulsion force magnitudes to be positive.

When we consider an edge that is part of a boundary, we can allow edges to cross because we now have a clear way to orient the interaction. The inside of the cell can be taken as a negative position with respect to the normal vector that points to the outside of the cell. To take advantage of this, we need to pay more attention to how we define the unit normal vector.

We can define a unit tangent vector **v** pointing along the length of the edge, and with this we can construct a unit normal vector **u**, by noting that for two vectors in the (*x, y*) plane, 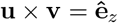. This is a unit normal pointing upwards out of the plane when viewed from above. Since by definition they are perpendicular, **u** and **v** are related by

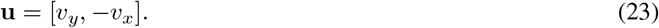

As a result, we define the interior of the cell to be the domain to the left when traversing the boundary of the cell in an anti-clockwise direction. The force is then

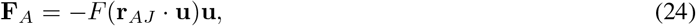

where the dot product will evaluate to negative values when *J* is inside the cell.

#### Applying a force to endpoints

As described above in *Finding interactions*, the point of action is defined as the first point an expanding circle touches a given edge. Around an endpoint, this produces a curved region of interaction with a constant radius. Figure 7 shows the interaction region around an endpoint that is part of a cell boundary. Likewise, Figure 6A seen previously shows the curved region around the endpoints of an isolated edge. If a node falls into this part of the interaction region, the interaction force is applied directly to the endpoint. When this happens, we can calculate the force on *A* in the same way as in Equation (22)

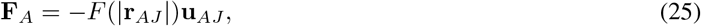

since, by definition, in this case A will lie on the endpoint.

**Figure 7:**
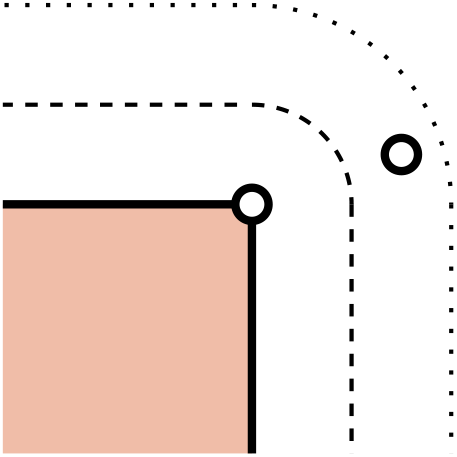
The interaction region around an endpoint that is part of a cell boundary.

The forces applied to a node are equal and opposite to the forces applied to an edge at the point of action. Therefore the force applied to a node due to the edge is

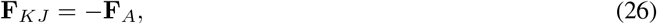

where *K* is an edge that *J* is not part of, and *A* is the point of action on *K* due to this node-edge interaction. Depending on the situation **F**_*A*_ may be defined by Equation (22) or by Equation (24). This also applies to the opposite node in a node-node interaction calculated in Equation (25).

In the models presented here, we resolve forces applied to a node without considering the impact to any edge it is part of. The nodes (endpoints) are treated as free objects in space, fitting in with the *semi-rigid body* concept arising from the edge force. For a given node *J*, the total force is found simply by summing all directly applied forces

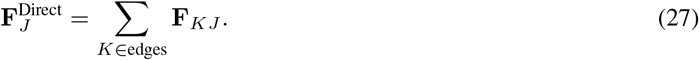

where *K* is any edge that does not have *J* as an endpoint.

#### Internal cell forces

At the individual level, we need to consider the forces required for a cell to maintain its structure. For a cell represented by an isolated edge, this means we need to control its length. We can do this by treating the edge as spring with a certain natural length and a force law acting to push the endpoints to the preferred separation. In this case we use a simple linear spring force

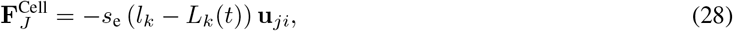

where the negative sign ensures the endpoints are pushed together when they are too far apart. Here **u**_*ji*_ is the unit vector pointing from endpoint *j* to endpoint *i*. The force law is a function of the difference between current measured length *l*_*k*_ and target length *L*_*k*_(*t*), and is parameterised by *s*_e_, which controls the magnitude of the force.

Polygonal cells require mechanisms based on its measured and target shape to represent the internal forces that give the cell structure. Here we use energy methods to drive the area and perimeter to specified values in a similar manner as used by Nagai and Honda [20] for the vertex model. The energy embodied in cell M is calculated by

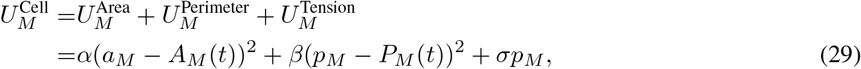

where *a*_*M*_ represents the current area, and *A*_*M*_ (*t*) the target area (which changes based on the cells age), and where *p*_*M*_, and likewise *P*_*M*_ (*t*), represent the current perimeter and its target value. The parameters *α* and *β* are energy density factors controlling how strongly the cell wants to return to its preferred area and perimeter, while *σ* is the total energy density of the surface due to surface tension. In a standard vertex model, the surface energy will depend on the relative affinity of the cell to bond with other cells and its environment. For a polygon cell model this would mean each edge has a value of *σ* determined by its immediate neighbourhood. To properly implement this, we would need to consider edge-edge interactions which we leave for later work. For the purpose of these illustrative examples, we make the simplifying assumption that any given edge will not have a relative preference to bond with anything in its neighbourhood, hence *σ* will be the same for all edges. As a result, the surface tension will tend to push the cell to have zero perimeter. This means the body of the cell will be under slight compression when all competing energies are at equilibrium.

The forces generated by the embodied energy, as with the vertex model, are applied only at the endpoints; no internal forces are applied to the edges. For a given node *J*, the force due to the embodied energy of each cell *M* it forms part of, is calculated by

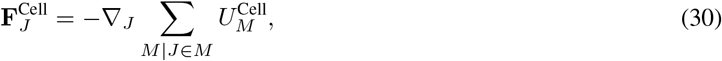

where ▽_*J*_ is the gradient with respect to the coordinates of node *J*.

For a polygon cell, its structure is controlled by both the target area and target perimeter as functions of time. These need some degree of synchronisation; more material inside the cell, will put strain on the surface membrane unless it also increases the amount of material making up the surface. To reflect this fact, the target perimeter *P*_*M*_ (*t*) is driven by the target area. For a general polygonal cell, we assume it wants to maintain the same perimeter as a regular N-gon of area *A*_*M*_ (*t*), which can be expressed by the equation

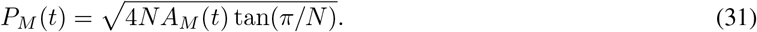

For a rectangular cell where growth is expected to happen primarily to the shorter edges (as is used in the epithelial monolayer model), the perimeter is determined by

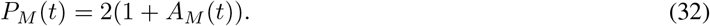

#### Cell Growth and Division

From a mechanical perspective, the function of the cell cycle is to control the growth of a cell. We implement growth by setting a target size, and letting the internal cell forces (i.e. 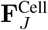) expand the size of the cell. At the beginning of the cell cycle, we assume the target size of the cell remains constant until it reaches age *t*_0_. After a cell has reached this age, it is permitted to start growing as long as its measured size is at least *s* > *γS*_0_, that is, the measured cell size (length or area) must be greater than some fraction of the resting target size. The parameter *γ* is the contact inhibition fraction, and will usually be 0 < *γ* < 1. If the size requirement is not met, the cell remains in a contact inhibited state until its measured size is large enough to commence growth. Growth occurs over an additional period of time t_*g*_, where the target size increases linearly from *S*_0_ to *S*_grown_. Division occurs as soon as growth is finished regardless of the measured size. The total time between cell birth and division is given by *t*_cycle_ = *t*_0_ + *t*_CI_ + *t*_*g*_, where *t*_CI_ is the unknown period of time spent waiting in a contact inhibited state. Depending on the state of the simulation, cells may remain contact inhibited indefinitely, and never start growing. To add stochasticity to the cell cycle and prevent adjacent cells from synchronising, the durations *t*_0_ and *t*_*g*_ can be chosen from a random distribution. When a cell reaches the end of its cell cycle, it divides according to the specific division mechanism for the cell type in the model.

#### Solving the equations of motion

Combining all of this together to calculate the motion of a node, we sum the directly applied forces, the rotation forces due to the edges, and the internal cell forces to give

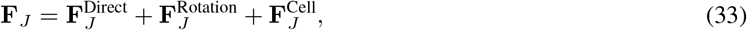

then apply the Forward Euler method to find its new position

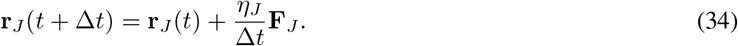

#### Practical model implementation details

Beyond the calculation of forces that constitute the intended behaviour of the model, there are practical decisions that need to be made in the modelling process and the implementation of the numerical library. These can be substantial additions in order to stop unphysical situations, or small details that nevertheless can affect the tissue level behaviour observed.

Isolated edges require special care to make sure the simulation remains physical (i.e. by stopping edges from crossing). Edges that are part of a polygon cells are no different. If an edge around a cell boundary is small, and the angle it makes with connected edges is also small, we can end up with edge inversion, where the boundary forms a non-simple polygon as shown in Figure 8. Small edges can also cause issues in applying the division process even with simple polygons. One way to prevent edge inversion is to use a force based on the length of the edge as done in Equation (28). This can be a simple linear spring that forces edges to a preferred natural length. When dealing with polygonal cells that can grow in size and divide, it is helpful to consider how edges contribute to the whole boundary. Here, we implement a regularising linear spring force which is applied to all edges around the boundary in order to keep the edge lengths relatively consistent. For a node *J* that is part of edge *k*, both subsequently part of cell *M* which is made up of *N*_*M*_ edges, which has perimeter of length *p*_*M*_, we can define a regularising force as

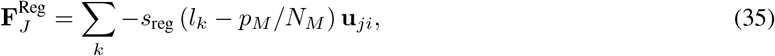

where *l*_*k*_ is its current measured length of edge *k* and **u**_*ji*_ is a unit vector from endpoint *j* to endpoint *i*. In words, each edge around a cell experiences a force so that they all tend toward the same length, being an equal fraction of the current measured perimeter. We note here that this force has no physical basis, and it is used exclusively to equalise the lengths of the edges.

**Figure 8:**
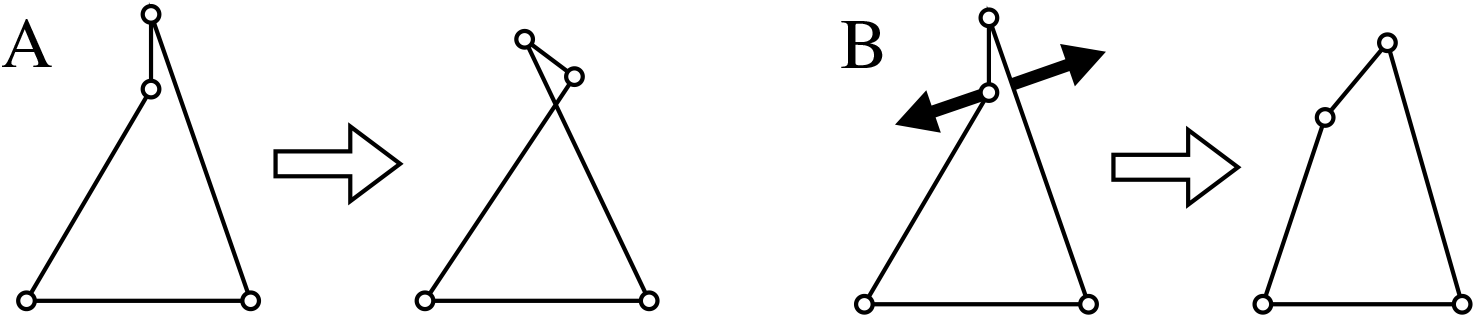
Edge inversion (A) is caused when small edges form small angles. In this situation, endpoints of an edge can approach closely to adjacent edges of the same cell, and without anything to stop them, they can cross in a subsequent time step. We can mitigate this by preventing small edges with a force to regularise the edge lengths. Small edges can be desirable in some instances, so we can also use the node-edge interaction mechanism to push edges of the same cell apart (B)

It is not always possible to prevent edges from becoming too small, even with the regularising force, and in fact sometimes we may want to allow small edges. To prevent inversion in these cases we target the formation of small angles by using the node-edge interaction mechanism to repel edges from the same cell. In order to invert, an endpoint needs to cross an edge from the same polygon, so we introduce a small repulsion-only node-edge interaction force to push them apart (Figure 8B). This will take the same form as Equation (21), but will only be applied when the node and edge are part of the same cell.

From a computational perspective, we can make some choices to speed up the simulation, or add in features to prevent undesirable tissue-level behaviours. As described above, the curved interaction regions mean that sometimes node-node interactions will occur. When standard neighbourhood searching algorithms are used, it is important to ensure that each interaction is only counted once. This needs extra care when searching within a single set of objects. For polygon cells, we find that the bulk of the interactions are in fact between node and edge. This situation means we are searching for interactions between two distinct sets, so each interaction can only be found once. Here we make the choice to neglect node-node interactions for all polygon cells to remove the need to validate interactions and speed up the neighbour-finding process. Practically, this can mean that nodes sometimes approach closer than the preferred separation distance. For non-connected polygon cells, this can on occasion be observed as a “reaching” behaviour, where nodes from adjacent cells reach out and come into contact. Motion of the nodes, however, rapidly brings them into range of a node-edge interaction, forcing them back into place. Neglecting node-node interactions seldom causes issues for connected polygon cells (such as those used in the epithelial layer model) since the angles between joined cells are usually greater than 130°, and often close to 180°. This means that most curved interaction regions are very small and reaching behaviour for connected polygon cells is almost never observed.

As with most node-based models where motion is the result of solving ODEs, there can be situations where forces get stuck in an unstable equilibrium. This can occur, for instance, where a row of nodes is is perfectly aligned with an axis and they experience compressive forces, also along that axis. In reality, a sufficiently large force will cause the row to buckle due to the fact that it is impossible for forces to be perfectly aligned. However, in the simulation, this unstable state can persist until rounding errors become large enough to tip the balance. To combat this, at the movement stage of the simulation, a small randomly directed force is applied to each node *J*:

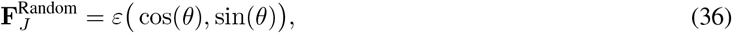

where 0 < *ε* ≪ 1, and **θ** is a randomly chosen angle, unique for each node at each time step. This small perturbation will kick the simulation out of unrealistic unstable equilibria, while being small enough to have no significant impact on the magnitude of the forces. Alternatively, this can be used to represent small random fluctuations caused by Brownian motion.

## Supporting information

Bacteria Video

Epithelial Video

Spheroid Video

## Code Availability

Code implementing the rigid body framework, along with all of the models presented here, and a guide for using the code can be found on https://github.com/luckyphill/EdgeBased

## Acknowledgements

PJB would like to acknowledge Janusz Krokiewski for his extremely thorough text on *Mechanics of a Rigid Body*, that was provided free of charge to students of mechanical engineering at The University of Melbourne, without which this work would not have been possible. This work was supported with supercomputing resources provided by the Phoenix HPC service at the University of Adelaide.

## Author information

### Affiliations

**School of Mathematical Sciences, The University of Adelaide, Adelaide,Australia**

Phillip J. Brown, J. Edward F. Green, Benjamin J. Binder

**School of Mathematics and Statistics, The University of Melbourne, Melbourne, Australia**

James M. Osborne

### Contributions

PJB conceived and developed the framework, developed the software, ran the simulations, and performed the analysis. All authors designed the exemplar models. PJB, JMO wrote the manuscript. JEFG, BJB and JMO supervised the project. All authors read and approved the final manuscript.

### Corresponding Author

Correspondence to James M. Osborne or Phillip J. Brown.

## Supplementary Methods

### Derivation of viscous rigid body mechanics

Rigid body motion in a system where inertia is negligible compared to viscous drag is a topic that is difficult to find in existing literature. As such, the equations of motion used in this work have been derived independently, following the method for inertial systems used in *Mechanics of a Rigid Body* by J. M. Krodkiewski [3].

To develop equations of motion for a overdamped/viscosity dominated system, we will assume firstly that all applied forces are balanced by drag forces, and secondly, that drag force is linearly dependent on velocity. The implicit assumption here is that the flow is laminar. This is sensible for the regimes we are concerned with in cell-based modelling where Reynold’s number is small. However it is worth noting that these equations of motion will not be valid for high Reynold’s number flow; it is a well established fact that drag in turbulent settings is quadratically dependent on velocity (*F*_*d*_ ∝ *v*^2^).

To start developing the over-damped equations, we look to the inertial equations for inspiration. Here we find several important components: total mass, centre of mass, angular momentum, and moment of inertia. To calculate rotation of rigid bodies in an over-damped setting, we need to establish analogous concepts based on drag rather than mass.

If we consider a two-dimensional body covering the region 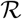, we make the definition

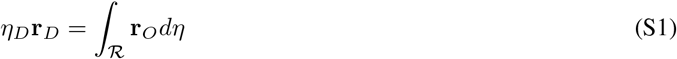

where **r**_*O*_ is the vector from the origin of the fixed system of coordinates to any point in 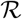, **r**_*D*_ is the *centre of drag*, and **η**_*D*_ is the *total drag* of the system. We intend the centre of drag to behave the same way as the centre of mass; it will be the point where we consider the total drag to be concentrated, and the point on the body where no rotation will occur if a force acts through it. Because of this, it is convenient to treat *D* as the origin of a coordinate system rigidly attached to the body, since this means we can separate out the translational and rotational motion components. We can then express any point on the body as the sum of the vector to *D* and the vector from *D* to the point in question, i.e.

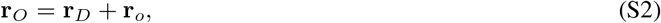

where we use *o* to represent vectors taken in the body system of coordinates. This allows us to rewrite the definition in Equation (S1) as

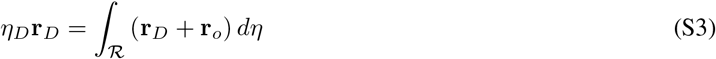

and since **r**_*D*_ is constant for the whole domain, this resolves to

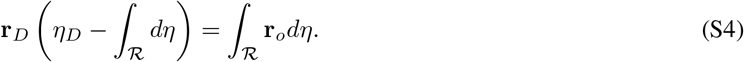

The right hand side is very similar to the definition of the centre of drag. It is the drag weighted averaged point taken over the body, except that the (arbitrary) origin starts at *D*. However, since we define the centre of drag to be precisely the point *D*, this means that

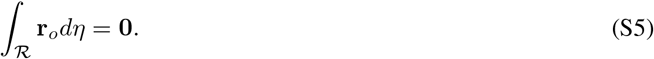

This leaves us with the condition

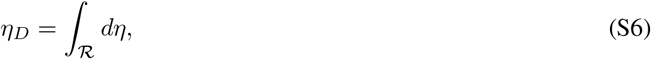

giving us a solution for the total drag. This takes the same form as calculating the total mass for an inertial body.

While this completely determines the total drag and centre of drag, it is not obvious that the centre of drag meets the requirements we set, i.e. if a force is applied at that point, the body does not rotate. To demonstrate this is the case, we consider the reactive forces on the body if we apply a force at some point *P*. The infinitesimal unit of force due to a unit of drag in the body is

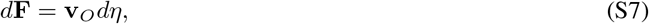

where **v**_*O*_ is the linear velocity of any point in the body with respect to the fixed system of coordinates. If we rigidly fix a set of coordinates on the body so that the origin passes through point *P*, and apply a force to that point, then we wish to find a constraint on *P* such that there is no resulting rotation about the origin. This means we require all of the moments about *P* due to the drag force sum to zero, i.e.

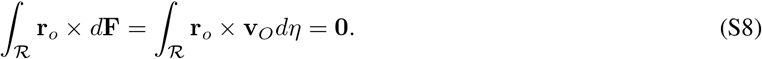

The vector **r**_*o*_ is the vector from *P* to any point in the body, but it is also the moment arm for the drag force due to each unit of drag in 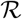. The velocity of any point in the body is derived from the position vector by taking the time derivative and noting that extra terms appear due to the possible rotation of the body. This results in

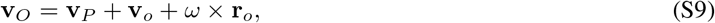

where **v**_*P*_ is the velocity of the body system of coordinates, *ω* is the angular velocity of the body (if any) and **v**_*o*_ is the velocity of any point in the body with respect to P. We know **v**_*o*_ must be zero since the axes are rigidly attached to the body, and we want the body to experience no rotation, so we set *ω* = **0**. Noting that **v**_*P*_ does not vary across the body, this results in

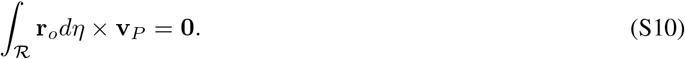

The velocity **v**_*P*_ cannot be zero since we are applying a non-zero force, leaving us with precisely the expression derived in Equation (S5). If the point *P* is not the centre of drag, then computing the integral will produce a vector that points from *P* to *D*. But the equation requires this vector to be zero, hence *P* = *D*, i.e. the point where an applied force causes no rotation must be the centre of drag.

To develop a concept equivalent to angular momentum, we consider an infinitesimal unit of the body 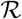 with drag *d*η** rotating about the origin *o* at a distance **r**_*o*_ with angular velocity *ω*. Its instantaneous linear velocity is

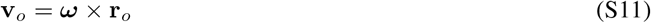

which will result in an infinitesimal viscous drag force

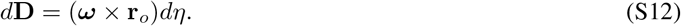

Since the drag force at each point is acting at a distance from the centre of rotation, it produces a torque on the body

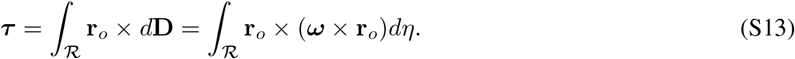

Applying the triple cross product and noting that we are working in two dimensions, this resolves to

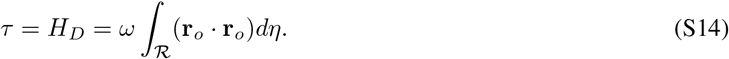

We call this torque the *angular drag*, *H*_*D*_ of a body, analogous to the angular momentum. This result may be counter-intuitive for those familiar with inertial systems, since the angular momentum (sometimes called the moment of momentum) is a distinct concept from torque. However, we remind the reader that motion is resisted by drag forces and not by momentum/inertia, hence it is sensible that we arrive at a “moment of drag force” i.e. a torque. The angular drag of the body must balance all of the applied moments, hence we can state that

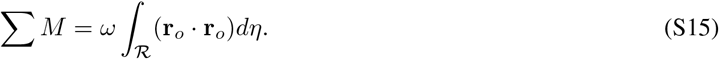

The integral quantity takes the same form as the moment of inertia in an inertial system, hence we name this the *moment of drag* and give it the symbol *I*_*D*_. Combining this with the linear motion of the body gives us the equations of motion for rigid body where drag is the dominant factor:

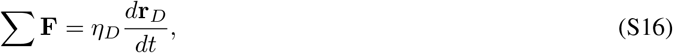

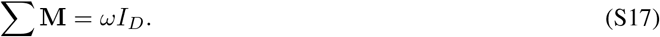

That is, the sum of the forces applied to the centre of drag is balanced by the velocity of the centre of drag, multiplied by the total drag, and the sum of the applied moments is balanced by the moment of drag taken about the centre of drag, multiplied by the body’s angular velocity.

## Supplementary Discussion

### Computational considerations and performance

In order for the edge based approach to function, it is necessary to consider precisely how neighbouring node-edge pairs are found. In an over-lapping spheres model, this can be done efficiently using a space partitioning method, whereby the spatial domain is broken up into “boxes” [1]. Each node is placed into its respective box based on its position, with the box being updated as the node moves through space. This makes finding neighbours much faster, as only the given box and its surrounding boxes need to be searched, rather than the entire simulation domain. A similar approach can be used for edges, where an edge is placed into each box that it passes through. Neighbourhood searching like this can be done in *O*(*kn*) time, where *n* is the number of objects, and *k* is the average number of near-neighbours [1]. As such, models using the rigid body framework should be able to obtain a similar time complexity to node-based approaches. This, however, will depend on how other components of the simulation are implemented (such as the force laws), which can become more involved for more complex cell constructions.

**Table S1:** Time taken to complete 40,000 time steps for each model presented, along with the starting and ending number of model components. Simulations were performed on a single core of the *phoenix* supercomputing cluster provided by The University of Adelaide. For the over-lapping rods model, one time step is 0.005 minutes, while for the spheroid and epithelial ring models, one time step is 0.005 hours, making for a total of 200 minutes and 200 hours respectively.

| Model |  | Nodes | Edges | Cells | Running Time |
| --- | --- | --- | --- | --- | --- |
| Over-lapping rods | start | 40 | 20 | 20 |  |
|  | end (avg) | 9554 | 4777 | 4777 | ~27hrs |
| Tumour Spheroid | start | 40 | 40 | 4 |  |
|  | end (avg) | 6280 | 6280 | 628 | ~17hrs |
| Epithelial ring | start | 40 | 60 | 20 |  |
|  | end | 3500 | 5250 | 1750 | ~12.5hrs |

Table S1 lists the running times for the three models presented here. Each instance of the models were run for 40,000 time steps (equivalent to 200 simulation hours or 200 simulation minutes in the case of the *E. coli* model) on a single core of the HPC cluster *phoenix* provided by the University of Adelaide (2.4GHz with 8GB RAM). Without a thorough a computational comparison controlling for hardware, software and implementation, it is difficult to quantitatively measure the performance differences between the models presented here and other existing modelling approaches. Additionally, each of the three model types has a unique relationship between numbers of cells, edges and nodes, not to mention the rate of new cell production depending on contact inhibition. Nevertheless, we can make some broad-brush assertions about the performance of the rigid body framework.

Between the three models, there is a positive correlation between running time and number of nodes, while there appears to be no correlation with number of edges. This is likely because, whilst we have added edges to the simulation, the neighbour finding algorithm is driven by nodes. When searching for a neighbour, we traverse the list of nodes and search their surrounding space for edges. There will be a natural limit to the number of edges that can be in proximity to a given node due to the configuration of the simulation, meaning that the *k* in the time complexity representation *O*(*kn*) should not vary wildly between model types. We also do not search for edge-edge interactions, meaning that n will only be the number of nodes. Therefore we can be reasonably confident that (at least as far as these models are concerned) a rigid body framework will be at no significant disadvantage to a node-based approach that relies on neighbourhood searching.

The augmented vertex model used for the epithelial ring should be able to run in a similar time frame to the standard vertex model when the tissue is not likely to self-contact. As mentioned above, the neighbourhood searching algorithm operates in *O*(*kn*) time, which evidently tends to zero when there are few neighbours. When collision is likely, this will increase roughly proportionally to the number of nodes in proximity of a contact point. Depending on the tissue implemented, this could tend towards the same complexity as a polygon cell model as the external surfaces become uniformly contacted, as observed in the epithelial model.

Of the existing deformable, non-connected cell methods, the polygonal cell model is perhaps closest to the immersed boundary method [4]. In this approach, cells have numerous nodes around their boundary (images from [4] show 58 in one case), and cells interact via node-to-nodes forces. Motion is determined by simulating a fluid flow, which then influences the movement of the boundary nodes. This gives a high level of detail for the shape and interactions, (perhaps capturing the level of individual adhesion molecules) and places the underlying physical mechanisms on very strong foundations. Evidently, though, this comes at a computational cost. The increased number of nodes, coupled with the need to solve, then interpolate the Navier-Stokes equations, makes a single time step take many times longer than for the polygonal cell model. An immersed boundary tumour spheroid simulation where cells had a cycle of roughly 5 simulation hours took “a few hours” on a single processor (2.7GHz) personal computer to reach a cluster of 50 cells [4]. For the decagon cell model, cells with a cycle length on average 20 hours reached a cluster size of 50 after about 15 minutes on a 2017 MacBook Pro with 2.3GHz processor. This is in no small part due to the ability to model cell boundaries with many times fewer components.

Conversely, to take proper advantage of the ability to represent arbitrary cell shapes, 6 or more nodes and edges is a reasonable minimum. Evidently, this means the simulation will take at least 6 times as long as a comparable overlapping spheres simulation, and at least twice as long as a vertex model simulation, where each node is usually part of three cells. This method is therefore best suited to intermediate scales, where the shape of cells are of interest, but the maximum number of cells is not too large.

While the rigid body framework can be used in a wide variety of situations, it is not suitable in every case. When used to model individual cells, as in the polygonal cell model, it can become computationally intractable for very large number of cells (several thousand). This can be mitigated by parallelisation, but it won’t be able to compete with a single node cell-centre model, which can handle in the hundreds of thousands of cells [2].

Overall, in the trade off between model detail, computational time, and implementation difficulty, a modelling paradigm using the rigid body framework would sit somewhere between the simpler node-based models (over-lapping spheres, tesselation, vertex) and the more detailed options (subcellular element, and immersed boundary). It trades the need for handling vertex swaps for the need to find neighbours and calculate rotations, but by doing this gains free motion of polygon shaped cells, and interactions between surfaces. It can capture the deformability and free shape afforded by the subcellular element and immersed boundary methods (albeit in a simplified form), while significantly cutting down on the number of simulation objects needed. This makes it potentially very useful in the niche between highly detailed small tissue simulations, and large scale tissues with simplified cell approximations.

## Supplementary Videos

### Bacteria video

This video demonstrates the over-lapping rods model replicating the experiment due to Volfson et al. [5] for a single random seed. Initially 20 randomly oriented cells are placed in the centre of the channel. The colour of a given cell indicates its angle from horizontal, from yellow (horizontal) to blue (vertical). Cells start to grow in localised clusters with roughly the same orientation. As the clusters grow and merge, the cells interact through the node-edge interaction mechanism, causing them to smoothly move and reorient themselves. As the proliferating front travels down the channel, the walls influence the orientation of the new cells, keeping them largely horizontal.

### Spheroid video

This video shows the polygon cell model producing a tumour spheroid. Four decagon cells are placed in the centre of a plane and left to proliferate. As the cell population expands, cells towards the centre become too compressed to start growing, leaving a narrow band of proliferating cells around the outer radius. Occasionally, inner cells start growing due to local pressure relief.

### Epithelial monolayer video

This video shows the progression of an unconstrained epithelial ring. The ring starts off being perfectly circular with equal size cells. As it grows, local areas of high pressure cause the ring to buckle. After enough buckling, the ring starts to contact itself in numerous places. The node-edge interaction mechanism along with the viscous rigid body laws of motion, allow contact forces to be transferred across the contact points, causing dynamic restructuring of the layer seen as secondary buckling.

## Footnotes

2 Some notable exceptions include Merks et al. [18] who use random movement to reduce total energy via metropolis sampling, and Smith et al. [30] who use movements that resolve overlaps in cells.

3 We are able to superimpose these movements because the linear movement represents the new position of the centre of drag, while the rotational movement expresses how the body is oriented. The motion can be split up this way because a force applied at a distance from the centre of drag can be represented as a force at the centre of drag, plus a moment [17]

